# Graph Contrastive Learning of Subcellular-resolution Spatial Transcriptomics Improves Cell Type Annotation and Reveals Critical Molecular Pathways

**DOI:** 10.1101/2024.03.08.584192

**Authors:** Qiaolin Lu, Jiayuan Ding, Lingxiao Li, Yi Chang, Jiliang Tang, Xiaojie Qiu

## Abstract

Imaging based spatial transcriptomics (iST), such as MERFISH, CosMx SMI, and Xenium, quantify gene expression level across cells in space, but more importantly, they directly reveal the subcellular distribution of RNA transcripts at the single-molecule resolution. The subcellular localization of RNA molecules plays a crucial role in the compartmentalization-dependent regulation of genes within individual cells. Understanding the intracellular spatial distribution of RNA for a particular cell type thus not only improves the characterization of cell identity but also is of paramount importance in elucidating unique subcellular regulatory mechanisms specific to the cell type. However, current cell type annotation approaches of iST primarily utilize gene expression information while neglecting the spatial distribution of RNAs within cells. In this work, we introduce a semi-supervised graph contrastive learning method called Focus, the first method, to the best of our knowledge, that explicitly models RNA’s subcellular distribution and community to improve cell type annotation. Focus first constructs gene neighborhood networks based on the subcellular colocalization relationship of RNA transcripts. Next, the subcellular graph of each cell can be augmented by adding important edges and nodes or removing trivial edges and nodes. Focus then aims to maximize the similarity between positive pairs from two augmented views of the same cell and minimize the similarity between negative pairs from different cells within a common batch. Guided by a limited amount of labeled data, Focus is capable of assigning cell type identities for the entire datasets at high accuracy. Extensive experiments demonstrate the effectiveness of Focus compared to existing state-of-the-art approaches across a range of spatial transcriptomics platforms and biological systems. Furthermore, Focus enjoys the advantages of revealing intricate cell type-specific subcellular spatial gene patterns and providing interpretable subcellular gene analysis, such as defining the gene importance score. Importantly, with the importance score, Focus identifies genes harboring strong relevance to cell type-specific pathways, indicating its potential in uncovering novel regulatory programs across numerous biological systems. Focus is freely accessible at https://github.com/OmicsML/focus.

## 1 Introduction

Recent developments in imaging-based spatial transcriptomics (iST) [28, 35, 14, 41, 10, 1, 5, 18] empower us to measure individual RNA molecules at subcellular resolution, which, in turn, facilitates the revelation of intricate spatial distribution of distinct cell types, and more importantly the cell-type specific subcellular localization of the transcripts. Consequently, these technologies hold exciting potential to shed light on the gene expression dynamics and regulation across many cell types at unprecedented spatial resolution. However, analyzing such data to achieve these goals will introduce several unfathomable computational hurdles. Among them, cell type annotation stands out as a particularly formidable challenge due to its low number of quantified genes, which results from the intrinsic limitation of the diffraction limit of light, the spectral overlap of the fluorescence probes, etc. For example, the Nanostring CosMx [14] platform defines the subcellular expression of merely 960 genes. Similarly, Xenium from 10x Genomics [18] measures only hundreds of genes in total. While MERFISH [5] in principle can measure the entire transcripts through a sophisticated multiplexing scheme, it usually can only robustly measure a few hundred genes.

Earlier methods on cell type annotation mostly rely merely on gene expression values but not the spatial localization of each RNA molecule in a cell. In more detail, single cells are first clustered by these methods, which are then manually annotated by experts based on cluster-specific markers [3, 12, 13]. Such manual annotation method requires substantial domain knowledge, and is time-consuming and laborious, making it less scalable to large and complex datasets. Subsequently, machine learning and deep learning techniques [25, 8, 4, 23, 31, 38, 32] have been employed to tackle these challenges. Among them, graph-based methods [31, 38, 32] stand out as a natural model paradigm of spatial transcriptomics by representing cells and genes as interconnected graphs, while also allowing seamless incorporation of prior knowledge. Furthermore, the utilization of advanced Graph Neural Networks (GNN) [20] is pivotal in capturing non-linear topological relationships among cells, potentially enhancing their performance in the task of cell type annotation. However, those methods often overlook the intracellular spatial distribution of RNA transcripts at the single molecule level, which potentially plays an important role in elucidating unique cell-type specific subcellular regulatory mechanisms. To incorporate spatial information of transcripts and their subcellular communities into cell type annotation, we propose Focus, a semi-supervised Graph Contrastive Learning (GCL) based algorithm of iST at subcellular resolution (Figure 1). To model the subcellular distribution and functional interaction between RNAs, Focus constructs gene neighborhood networks that constitute the basis for all downstream modeling. To learn subcellular features for each cell, we next employ GCL framework on the gene neighborhood network [7, 2, 40] by maximizing the similarity between positive pairs from the same cell and minimizing the similarity between negative pairs from distinct cells within a common batch where the graph pairs are generated via augmented graphs through adding or removing graph nodes or edges. After the GCL process, Focus computes an important score for every transcript and RNA subcellular community (the subnetwork formed by a subset of RNA transcripts), reflecting their importance for that cell. We find the genes with high scores tend to have significant importance and relevance to specific cell types based on detailed gene set enrichment analyses. We further demonstrate the effectiveness and robustness of Focus across various platforms, including CosMx, MERFISH, and Xenium, and different numbers of cell types (ranging from 8 to 24) in the context of cell type annotation.

**Figure 1:**
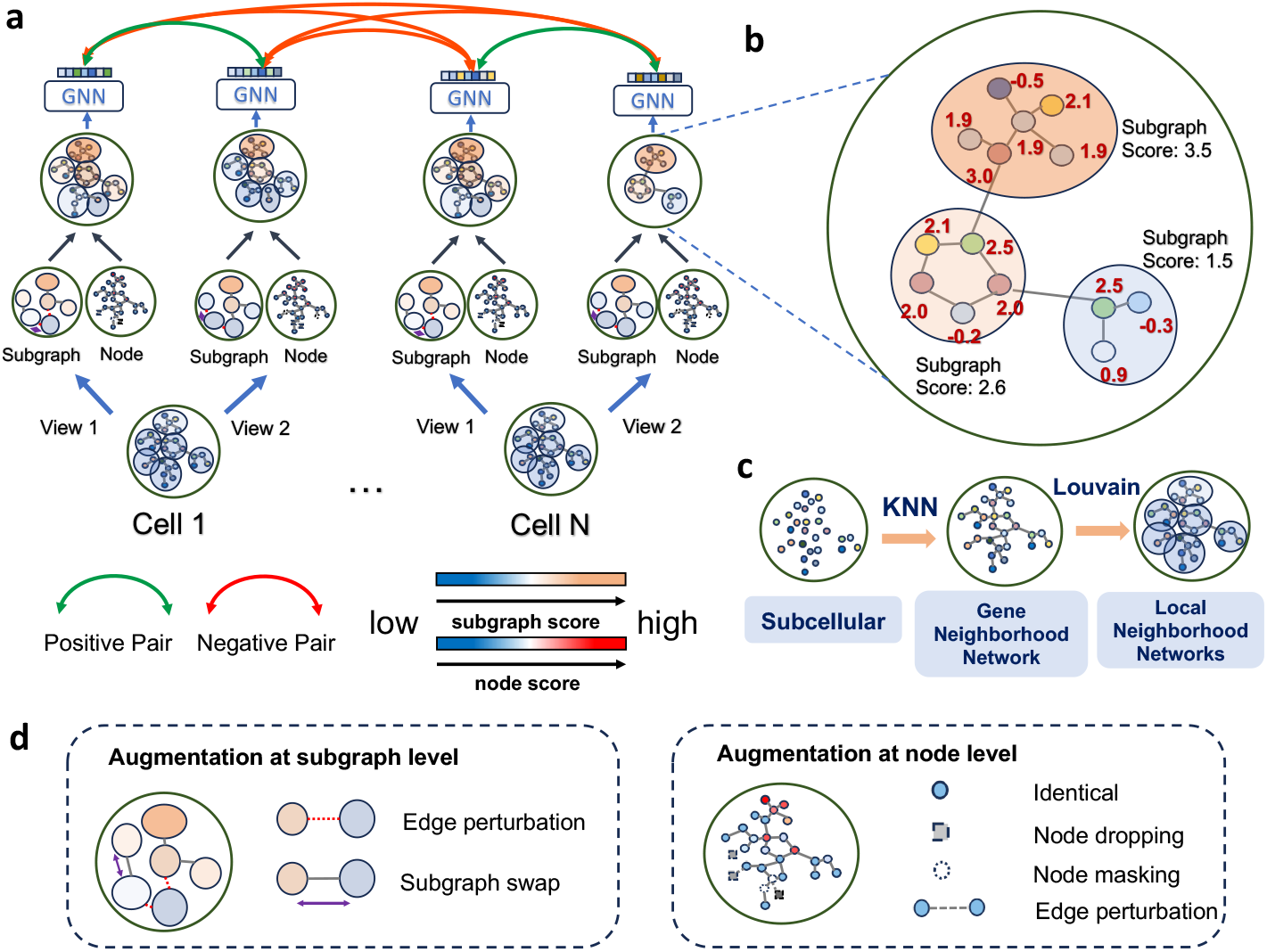
An Overview of the Focus framework. **a**. The graph contrastive learning workflow in Focus. **b**. Learned scores for each node and subgraph after graph contrastive learning. **c**. Gene neighborhood network construction from subcellular information. **d**. Augmentation methods at subgraph and node level.

Focus demonstrates significant improvements over state-of-the-art algorithms. In particular, using the same number of labeled data, i.e., annotated cells, as the baseline, Focus demonstrates superior performance on all test datasets, achieving improvements up to **10%** in terms of accuracy and **40%** in terms of F1-score. It is important to note that our method exhibits high robustness, performing relatively well even when provided with a limited amount of labeled data. In contrast, baseline methods show a rapid decline in performance as the amount of labeled data decreases. In addition, we rank genes based on the importance score from Focus and subject genes with the higher scores to gene sets enrichment analysis and find that they are highly enriched in cell type-specific pathways. Our main contributions in this paper are:

- We introduce Focus, a GCL model to enhance cell type annotation on spatial transcriptomics data by leveraging subcellular spatially resolved gene neighborhood networks. To the best of our knowledge, this is the first work that uses the spatial information of the transcript and the RNA subcellular communities in the cell to improve cell type annotation.
- Focus exhibits high robustness even when provided with a limited amount of labeled data, and outperforms state-of-the-art methods by a large margin across various iST platforms.
- Based on learned importance scores for genes by Focus, we identified novel genes with higher scores that are highly relevant to cell type-specific pathways.

## 2 Related Work

In the realm of single-cell analyses, one of the essential objectives is to discern and categorize the diverse cell types within a mixed population of cells, making cell type annotation a pivotal undertaking for researchers. The task on iST presents a unique challenge due to its general low number of measured genes or often high dropout rate. Presently, the majority of cell-type annotation methods are tailored for single-cell RNA seq data. CellTypist [8] builds on a logistic regression model with stochastic gradient descent learning for cell type identification using a curated pan-tissue database for immune cell types. TOSICA [4] is a multi-head self-attention network for interpretable cell type annotation in single-cell data by connecting attention to prior biological knowledge such as pathways or regulons. ACTTINN [23] trains a neural network using reference datasets to predict cell type for each cell. scDeepSort [31] and sigGCN [38] are two methods that draw upon graphs. sigGCN employs a graph convolutional network (GCN) [20] to reflect the non-linear topological relationship among cells. It first constructs a gene-wise weighted adjacency matrix using the STRING database [34] to create a gene interaction network where node features are defined as corresponding gene expressions. This graph is used as the input to a GCN-based autoencoder. The training objective consists of the gene expression reconstruction loss, the classification loss, and finally a regularization loss to prevent overfitting. scDeepSort uses a Graph Neural Network (GNN) on a weighted bipartite graph where both cells and genes are its nodes. The gene expression value for each cell-gene pair is used as the edge weights. After graph aggregation, cell node representations in the latent representations produced by the aggregation layer pass a linear classifier to predict cell labels. spatial-ID [32] is one of the few methods designed for spatial transcriptomics. It is also a GCN-based method where an autoencoder [15] is used to encode gene expression profiles and a variational graph autoencoder [21] is used to embed spatial information of cells simultaneously. spatial-ID takes into account the spatial information of cells but does not yet consider the spatial information of the RNA transcripts. TACCO [24] is also an approach designed for cell type annotation on spatial transcriptomics, and is an optimal transport-based method, enabling the transfer of annotations from single-cell data to spatial data.

## 3 Methods

The key idea of Focus is to involve subcellular localization of RNA molecules or RNA subcellular communities within cells to learn cell type-specific intracellular spatial distribution of RNA for cell type identification under the setting of limited or even no labeled data. We achieve the goal with graph contrastive learning. An overview of Focus is shown in Figure 1. The input of Focus is gene neighborhood networks constructed from transcripts within cells (Section 3.1). Each cell will produce two augmentation views, with each of both being generated by the combination of subgraph and node-level augmentations (Section 3.2). Given augmented graph views of each cell, a GNN model is employed to update node embedding and to learn subgraph embedding (Section 3.3). Focus aims to maximize the similarity between positive pairs from two augmented views of the same cell and minimize the similarity between negative pairs from different cells within a common batch guided by limited labeled data (Section 3.4).

### 3.1 Gene neighborhood network construction

Before performing gene neighborhood network construction, we process the original subcellular spatial transcriptomics data with the following steps:

- Elimination of low-quality cells: we exclude cells that are either non-viable or exhibit a transcript count below a predefined threshold denoted as *α*. The determination of the threshold *α* is dataset-specific; for the MERFISH, CosMx SMI, and Xenium datasets, values of 100, 50, and 50, respectively, are employed.
- Exclusion of negative control targets: given that negative control probes are designed to capture extraneous sequences not present in the tissue, serving as non-target controls to assess non-specific ISH probe hybridization, we filter out negative control targets and retain transcripts that corresponded to genes.

As shown in Figure 1c, given transcripts with spatial coordinates within each cell, we employ the KNN algorithm [9] to construct physical distance-based gene neighborhood networks where any transcripts will form a connection if the separation distance between them falls below a specific threshold *d*. Considering the diverse resolution in iST data, we designate values of 12, 2, and 3 for the parameter *d* in the context of CosMx SMI, MERFISH, and Xenium data, respectively. As a result, in the constructed gene neighborhood network, individual nodes represent single transcripts and are linked to an average of 10-30 neighbors. To better simulate transcript enrichment, a clustering method like Louvain [37] is employed to group the constructed graph into several local gene neighborhood networks, which refer to local RNA density or subcellular spatial regions to reveal the subcellular organization of RNA.

### 3.2 Data augmentation

Data augmentation is essentially the learning process of the intracellular spatial distribution of RNA transcripts for each cell type. Based on the constructed gene neighborhood network with subgraphs, augmentation can be performed at two levels (subgraph level and node level) as shown in Figure 1d:

- Augmentation at the subgraph level: **edge perturbation** involves the potential removal of edges between subgraphs, while **subgraph swapping** entails the exchange of two subgraphs.
- Augmentation at the node level: **identical** implies the preservation of edges or nodes in their original state, **node dropping** involves the removal of nodes, resulting in an alteration of the graph structure, **node masking** solely conceals node attributes while maintaining the graph structure, and **edge perturbation** pertains to the deletion of edges connecting nodes.

It is important to note that both subgraph and node level augmentation are automatically chosen for inclusion through learning in Focus, eliminating the need for manual selection of augmentation strategies. Please refer to more details in Section 3.3.

### 3.3 The proposed architecture

After graph construction introduced in Section 3.1, we have a gene neighborhood graph *G* = (*V, E*), where *V* =*{v*_1_, *v*_2_, …, *v*_*n*_*}* denotes a node set with *n* transcripts in the cell, and *E* denotes an edge set. Node feature is represented by *X* ∈ ℝ^*n×d*^ where the node embedding is initialized with one hot embedding with dimension *d*, and *d* corresponds to the total count of gene types. The graph structure is characterized as an adjacent matrix *A* ∈ *R*^*n×n*^. The graph would be further partitioned into *k* subgraphs *{***S**_**1**_, **S**_**2**_, …, **S**_**k**_*}* by a graph clustering algorithm like Louvain.

Focus consists of two convolutional layers to update node embeddings and subgraph embeddings. Specifically, each graph convolutional layer is defined as:

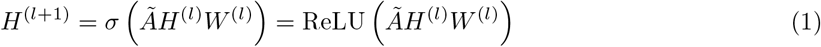

where 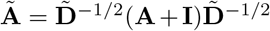 with 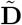 the diagonal matrix of **A** + **I** and **I** is the identity matrix. *H*^(*l*)^ is the input from the previous layer. *W* ^(*l*)^ is the weight matrix of the *l*-th layer. ReLU(·) is the nonlinear activation function. Here, the input for the first layer would be the original node representation H^(0)^ = X. The two convolutional layers in Focus are described as:

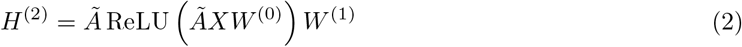

where *W* ^(0)^ and *W* ^(1)^ ∈ *R*^*d×d′*^ are weight matrices to be learned, and *H*^(2)^ is the learned node embedding after two layers of graph convolution.

Previous studies have revealed a strong correlation between the selection of specific components in a graph (such as nodes, subgraphs, etc.) and the intrinsic properties of the graph during contrastive learning. Consequently, the graph property can be effectively captured by retaining highly relevant nodes or subgraphs while eliminating less critical components [39]. Thus, a series of augmentation strategies for both subgraph and node levels are collected, including node dropping, node feature masking, edge perturbation, and subgraph swap. We employ the Gumbel-Softmax to obtain these strategies’ probabilities and set them as the score of nodes and subgraphs. These scores indicate the significance of these nodes or subgraphs within the overall graph or cell:

- **Node Level**: the score of the node is obtained by calculating the probability of node level augmentation, including node dropping, node masking, and edge perturbation within a subgraph. Thus, given a subgraph **S**, the score of node dropping can be formulated as:

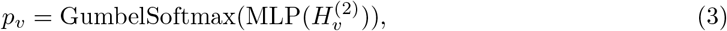

where MLP refers to a multilayer perceptron [27], 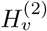 denotes the learned embedding of node *v* from GNN, *p*_*v*_ represents the score drawn from the distribution used to determine the nodes chosen for augmentation.
- **Subgraph Level**: the score of subgraph is obtained by calculating the probability of subgraph level augmentation, including subgraph swapping and edge perturbation between subgraphs. Thus, given a subgraph **S**, the score of the subgraph can be formulated as:

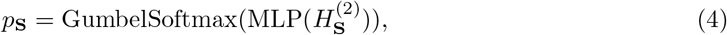

where 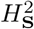 denotes learned subgraph embedding which aggregates from node embeddings within the subgraph, *p*_**S**_ is the score drawn from the distribution used to determine the subgraph chosen for augmentation.

#### Augmented Graph Assembler

For each partitioned subgraph *{***S**_1_, **S**_2_, …, **S**_*k*_*}*, node level augmentation and subgraph level augmentation will be performed separately. We denote the augmented view of **S**_*i*_ as 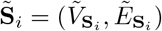. Thus the final augmented graph view 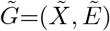 can be obtained via a combination of each subgraph as below:

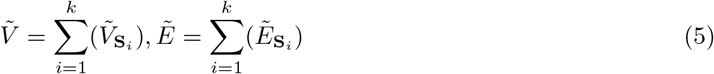

Upon completing the assembly of augmented subgraphs, we introduce the augmented graph into ResGCN [6] for the ongoing update of node embeddings and edge features. Ultimately, the graph embedding is obtained as 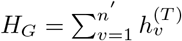, where *n′* represents the number of nodes in the augmented graph. Generally, Focus uses a Gumble-Softmax reparameterization trick to bridge the gap between subcellular graph augmentation and GNN back-propagation so that Focus has the capability to autonomously choose suitable nodes or subgraphs for augmentations.

### 3.4 Loss function

In order to better learn the property of the subcellular property of each cell, we follow the InfoMin principle [36] which states that good contrastive learning should maximize the label-related information while reducing the similarity between them. In the training phase, a data batch comprising N cells is randomly selected, and this batch is fed into the two graph view generators to generate 2N graph views. The two augmented views originating from the same input graph are treated as the positive view pair, while the views derived from distinct input graphs are considered the negative pair. In our work, we define contrastive learning loss ℒ_*cl*_ to maximize the uniformity between positive pairs and negative pairs. To be specific, we adopt the normalized temperature-scaled cross-entropy loss(NT-XEnt) ℒ_*cl*_, which can be formulated as:

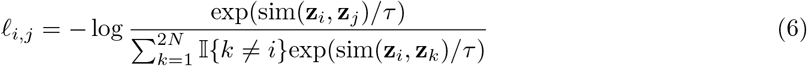

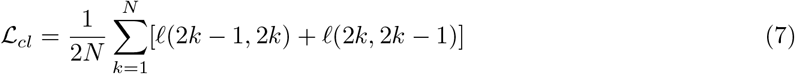

Here, (*i, j*) represents a positive pair of samples from the same cell, while (*i, k*) denotes a randomly sampled pair from the batch. **z**_*i*_ denotes the embedding of cell *i* obtained from the final output *H*_*G*_ of ResGCN. The cosine similarity between cell *i* and cell *k*, denoted as 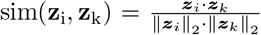, where *τ* represents the temperature parameter. The loss is computed collectively across all positive pairs within a batch. Different from other supervised learning methods that only train on reference labeled data and infer on query data, we design a novel semi-supervised setting that can effectively learn the inner property of query data with limited labeled data. Specifically, we use the whole test data without labels for unsupervised contrastive training and use limited reference data with labels to guide the model to learn cell type specific features. We then test Focus on query data. Here, we adopt the cross entropy loss ℒ_*cls*_ as supervised classification loss, which can be formulated as:

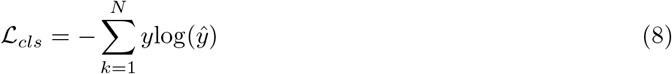

where *y* and *ŷ* denote ground truth labels and predicted labels, respectively. The overall objective ℒ of Focus can be formulated as:

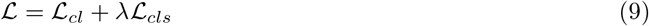

where *λ* is a hyperparameter to balance unsupervised contrastive learning loss ℒ_*cl*_ and supervised classification loss ℒ_*cls*_.

## 4 Results

### 4.1 Datasets and Evalutaion metrics

#### Datasets

To demonstrate the performance of Focus on cell type annotation, we collect a variety of subcellular spatial transcriptomic datasets from NanoString CosMx SMI [14], Xenium from 10x Genomics [18] and MERFISH [5] for evaluation. The tissues under evaluation encompass the human kidney, lung, breast, and mouse cortex.

##### CosMx Human Lung

This dataset is generated with a 960-plex CosMx RNA panel with CosMx SMI. There are 8 samples from 5 non-small cell lung cancer (NSCLC) tissues, including 3 female and 2 male patients. A total of 766, 313 cells are analyzed, with an average of 265 transcripts per cell, each represented by a subset of 960 selected gene targets. Each cell within the tissue is classified into one of 18 unique cell types.

##### CosMx Human Kidney

This dataset was generated using a 960-plex CosMx RNA panel with the CosMx SMI platform and collected from a kidney core biopsy taken from patients with lupus nephritis. We curated and selected 5 samples from 3 different patients, assigning 8 unique cell types as labels for the classification task.

##### MERFISH Mouse Primary Motor Cortex (MOp)

This dataset comprises spatially resolved gene expression profiles of individual cells within the mouse primary motor cortex, obtained through MERFISH. The dataset encompasses approximately 300, 000 cells and 258 genes, with an average of 614 transcripts detected per cell. Furthermore, it identifies a total of 24 distinct cell types within the mouse primary motor cortex.

##### Xenium DCIS

This dataset was created using Xenium from 10x Genomics and was collected from human breast tissue containing Ductal carcinoma in situ tumor cells. We conduct data cleaning and utilized two replicates from a single sample, resulting in the detection of approximately 300, 000 cells and 313 genes. The dataset encompasses a total of 19 distinct cell types.

#### Evaluation metrics

To assess the effectiveness of cell type annotation, we adopt the evaluation metrics *Accuracy* and *F1-score* following previous cell type annotation studies [8, 4, 23, 31]. We obtain the ground truth released from original datasets, and then calculate the *Accuracy* and *F1-score* between ground truth and predicted cell type labels generated by Focus or baselines. The *Accuracy* and *F1-score* are formulated as:

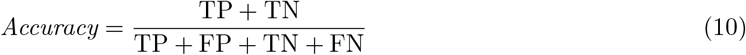

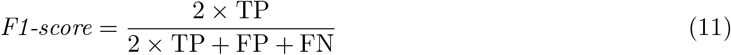

where true positive (TP) denotes the number of correctly classified positive samples; true negative (TN) represents the number of correctly classified negative samples; false positive (FP) indicates the number of samples incorrectly classified as positive, and false negative (FN) is the number of samples incorrectly classified as negative.

### 4.2 Focus achieves SOTA on cell type annotation across diverse iST platforms

To validate the efficacy of our Focus across diverse iST platforms, we perform validation experiments of cell type annotation on four datasets using Focus and state-of-the-art methods. These experiments encompass three iST platforms: CosMx, MERFISH and Xenium. The datasets cover a spectrum of cell type numbers and identities, ranging from 8 to 24 distinct cell types, and include both human and mouse samples, containing various tissues such as kidney, lung, and breast. In each dataset, we select one sample as the reference data and another as the query data. Please refer to the Supplementary informationB for other experimental settings. During the training of Focus, query data without label is also included and contributes informative data for contrasting and understanding the inherent characteristics of the query data. In practical scenarios, reference and query datasets may originate from various studies or different subjects, potentially introducing batch effects that can impair cell type annotation performance. Hence, we conduct our experiments under two conditions: utilizing samples from the same subject and samples from distinct subjects for comprehensive evaluation. We also benchmark scDeepSort[31], CellTypist[8], TOSICA[4], ACTINN[23], classical cell type annotation methods on single-cell RNA seq data and Tacco[24] which is specifically designed for spatially transcriptomic data on the above datasets for comparison.

As shown in Figure 2a and Figure 2b, we observe that regardless of whether the reference and query data are sourced from the same or distinct subjects, our Focus consistently outperforms state-of-the-art approaches both on accuracy and F1-score perspectives. Notably, on the Xenium DCIS dataset, Focus demonstrates a remarkable advantage, surpassing the second-ranked method by a substantial **10%** in terms of accuracy and **40%** in terms of F1-score. To be specific, Focus achieves 87.57% (accuracy) and 67.1% (F1-score) while the second-ranked method ACTINN gets 60.2% (accuracy) and 19.6% (F1-score) on Xenium DCIS dataset, The methods specialized in annotating single-cell data, such as CellTypist and scDeepSort, frequently employ the filtration of genes with exceedingly low expression levels in cells. Yet, this strategy can potentially constrain their capacity to harness the full limited gene information within spatial transcriptomics data. Tacco is specifically designed for spatial transcriptomics data, and it performs well in capturing cell-level spatial information on MERFISH and CosMx Lung datasets. However, when applied to other datasets featuring intricate cell types, it encounters notable variability in cell type prediction due to the complexity of cell type. Instead, our Focus focuses on acquiring cell type-specific information at the subcellular level, and this capability is not constrained by the quantity of cell types present. As a result, we consistently achieve top-tier performance, regardless of whether the dataset contains a limited or extensive variety of cell types.

**Figure 2:**
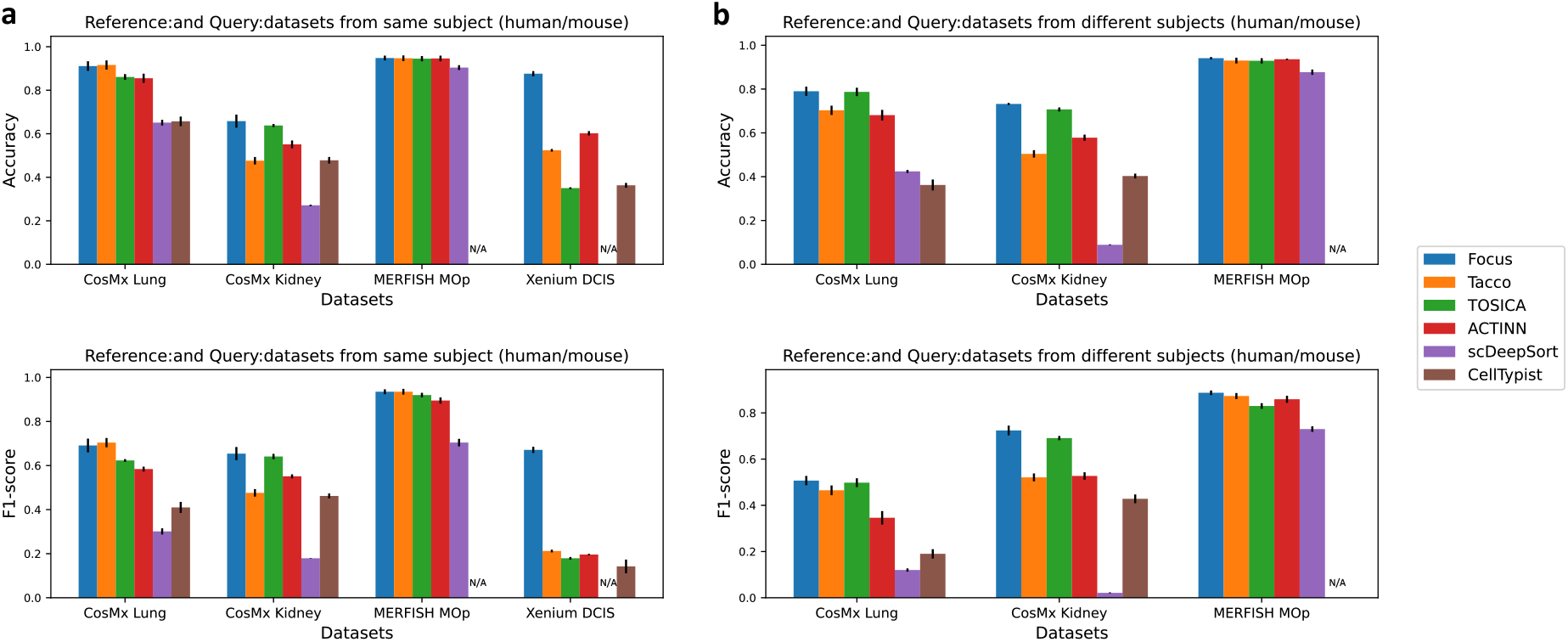
Benchmark of cell type annotation between Focus and state-of-the-art approaches. **a**. Reference data with label and query data from the same subject. **b**. Reference data with label and query data from the different subjects. *N/A* indicates that scDeepSort and CellTypist cannot work effectively on certain MERFISH MOp or Xenium DCIS datasets due to their limited capacity to process datasets with a low number of genes.

It is worth noting that on rather homogeneous datasets Focus still achieves superior performance (65.81% accuracy for the first setting) compared with other methods (scDeepSort: 27.1%, Tacco:47.6%, CellTypist:47.8%, TOSICA:63.8%, and ACTINN: 55.1%) on the CosMx kidney dataset. Among this dataset, there are three main cell types: proliferating proximal tubule cell, distinct proximal tubule 2 cell and distinct proximal tubule 1 cell, which are very similar to each other. This brings challenges to cell type annotation and these cell types are very difficult to distinguish by previous biological researches [22, 33], which mainly rely on gene expression values. The paramount aspect of Focus lies in the acquisition of cell type information from RNA’s subcellular spatial molecular pattern, which proves to be more effective when compared with methods relying solely on gene expression values. Furthermore, Focus presents remarkable transferability, even when applied to samples from distinct subjects. In contrast, other approaches experience a performance decline when confronted with reference data from different subjects. This heightened challenge in spatial transcriptomics data, characterized by a shallower read depth, presents difficulties in feature learning at the individual cell level. However, our model consistently attains the highest classification performance across all datasets.

### 4.3 Graph-based gene importance scores reveals enriched cell type-specific pathways

By leveraging the Gumbel-Softmax reparameterization technique for selective augmentations on each gene or transcript, we can derive scores for all genes, serving as indicators of their importance within data augmentation. To effectively illustrate the importance of each gene with respect to each cell type, we compile scores for all 960 genes, categorizing them by cell type within a dataset of 20, 645 sampled cells from CosMx Lung NSCLC. It is noteworthy to emphasize that these genes are not only associated with cell types but also play roles in various biological processes, including those related to hormones, hormone receptors, hormone processing, receptors, and ligands, as well as cell states, among others. Moreover, we carefully rank these genes according to their score distribution (based on the maximum). Based on the ranking and scores of these genes, we perform GSEA preranked gene set enrichment analysis using the gseapy package[11] and KEGG database[19].

We filter for pathways that are significantly enriched in the high-ranking genes judged by setting Normalized p-val (NOM p-value) *<* 0.05, False Positive Rate q-value (FDR) *<* 0.25, and Normalized Enrichment Score (NES) ≥ 1, as reported by previous work[17][29]. We find that those genes in the enriched pathways are mostly the ones with higher scores, and those pathways are highly relevant to the corresponding cell types. Given the fact that the CosMx dataset contains tumor cells and a large number of immune cells, as shown in Figure 3a, our enrichment analysis reveals that these cells predominantly exhibit functions highly related to antigen processing and presentation, ECM-receptor interaction, Epstein-Barr virus infection, which is consistent with their fundamental immunity functionalities. To better discern functional differences among different cell types, we select highly significant pathway information only in specific cell types as shown in Figure 3b. For NK cells, we have successfully observed highly significant signal pathways expressed within NK cells, such as natural killer cell mediated cytotoxicity. Similarly, we have identified signal pathways associated with tumor cells triggered by colorectal cancer, hypertrophic cardiomyopathy (HCM), and amyotrophic lateral sclerosis (ALS), which are consistent with previous research[26][30]. For B cells, we have discovered specific immunological pathways, such as primary immunodeficiency, an immune deficiency disorder primarily associated with B cell dysfunction[16]. These examples convincingly demonstrate that Focus can learn pathway information highly relevant to cell types, which plays a crucial role in cell type classification for spatial transcriptomics data.

**Figure 3:**
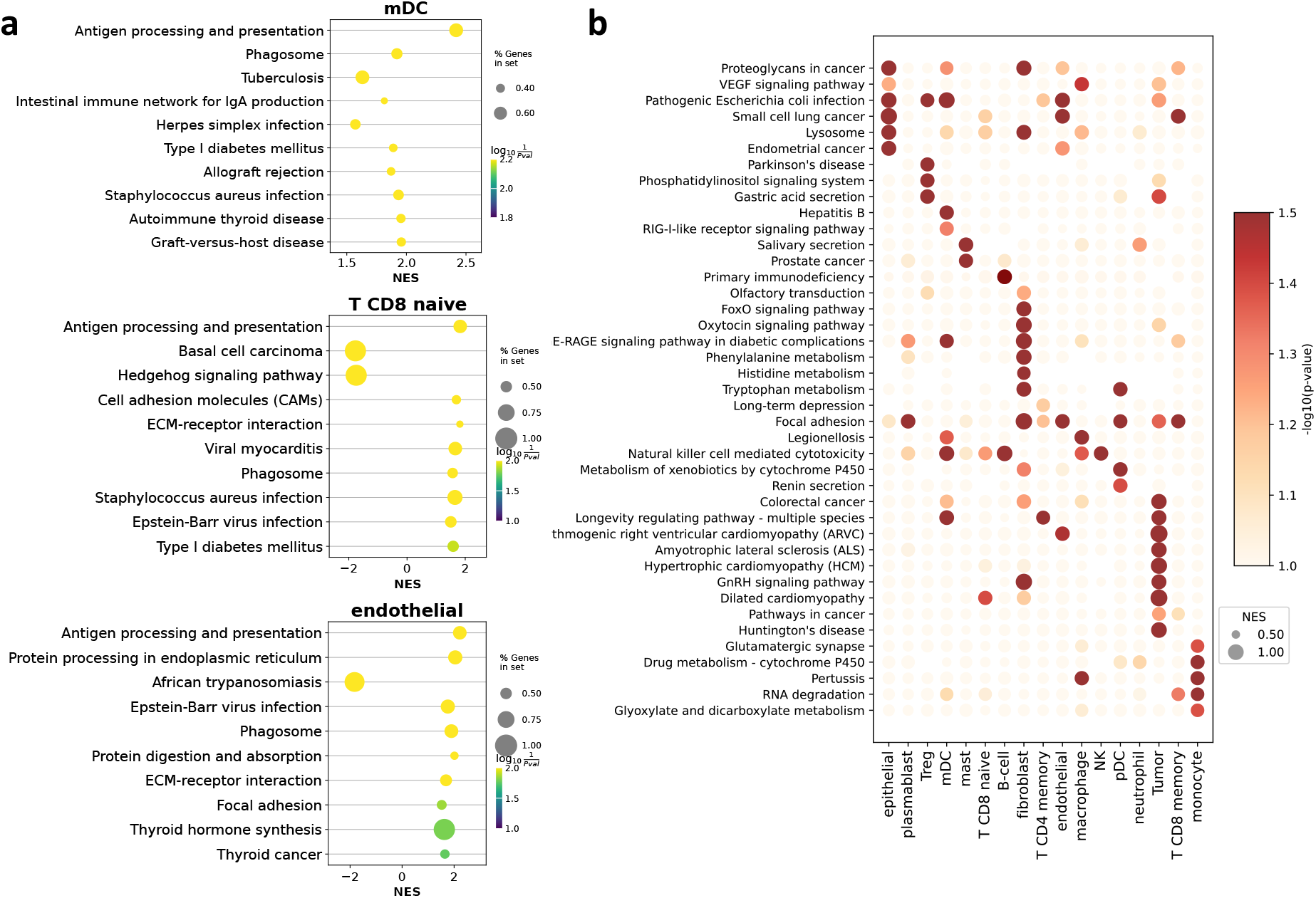
Enriched cell type-specific pathways based on gene important score from GSEA Preranked analysis. **a**. Enriched pathway of mDC, CD8+ naive T cell, and endothelial cell. **b**. Heatmap of selected cell-type specific pathway.

### 4.4 Focus is robust to limited reference data

Recognizing the challenge of acquiring extensive high-quality labeled reference data, we carry out experiments with a restricted pool of labeled reference data to assess the performance of each method. To achieve this, we devise three distinct scenarios where the query data remained constant at 9k cells, while varying the sizes of the reference data from 1k, 4k, to 9k cells. In order to ensure consistent findings and mitigate the influence of random variations, we systematically conduct a series of experiments across multiple samples: (1) Training on reference data CosMx Lung 5-3 and test on query data CosMx Lung 5-1 (Figure 4a). (2) Training on reference data CosMx Lung 5-3 and test on query data CosMx Lung 5-2 (Figure 4b). (3) Training on reference data CosMx Lung 5-1 and test on query data CosMx Lung 5-3 (Figure 5a in Supplementary A). (4) Training on reference data CosMx Lung 5-1 and test on query data CosMx Lung 5-2 (Figure 5b in Supplementary A). (5) Training on reference data CosMx Lung 5-2 and test on query data CosMx Lung 5-3 (Figure 5c in Supplementary A). (6) Training on reference data CosMx Lung 5-2 and test on query data CosMx Lung 5-1 (Figure 5d in Supplementary A).

**Figure 4:**
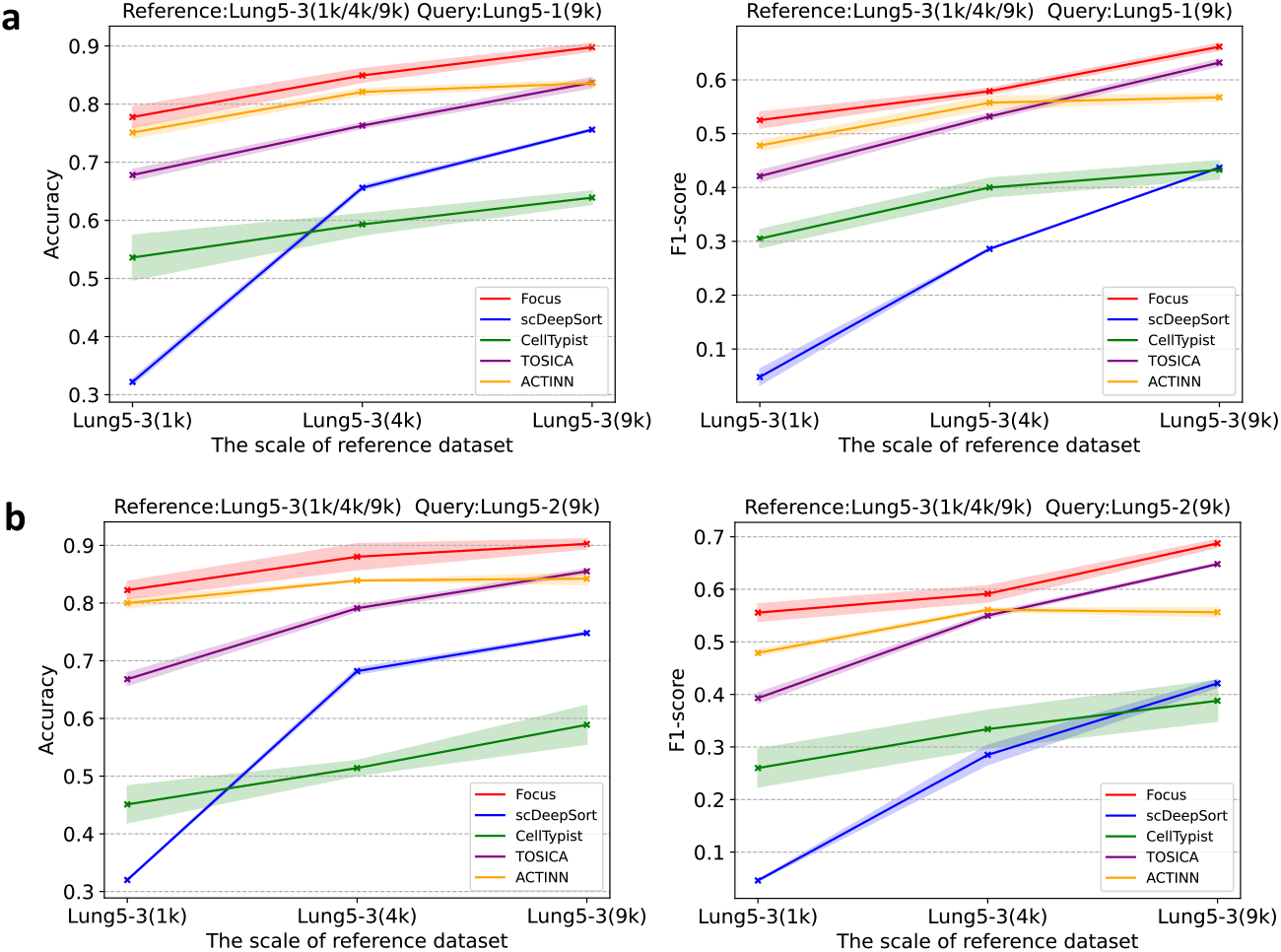
Robustness of Focus and other methods to the reference data size. **a**. Reference data: CosMx Lung5-3, Query data: CosMx Lung5-1. **b**. Reference data: CosMx Lung5-3, Query data: CosMx Lung5-2.

Our consistent observation is that, regardless of whether our Focus is trained with 9k, 4k, or even 1k reference data, the variation in reference data quantity has a minimal impact on query performance, and accuracy consistently remains above 80%. However, all other methods exhibit a performance decline as reference data decreases, albeit to varying degrees. For instance, scDeepSort experiences the most significant performance drop across all experiments, while cellTypist is less affected by data reduction but consistently demonstrates lower accuracy across reference data scales. ACTINN appears less affected overall. To sum up, our Focus maintains an 80% accuracy level across various experiments, demonstrating robustness to reference data downsampling from 9k to 1k, as well as to the variations in reference data, thus underscoring its robustness.

## 5 Discussion

Our Focus method for cell type annotation outperforms state-of-the-art approaches across various iST platforms, including CosMx, MERFISH, and Xenium, achieving substantial improvements (up to 10%) in accuracy, even when working with limited labeled data. Notably, Focus leverages subcellular spatial gene neighborhood networks, making it the first of its kind to utilize transcript’s subcellular and spatial community information for enhanced cell type annotation. Additionally, the learned importance scores from Focus reveal the relevance of highly scored genes to cell type-specific pathways, reinforcing the power of our approach in deciphering molecular mechanisms underlying intricate cellular processes.

