## supplemental file for "Graph Contrastive Learning of Subcellular-resolution Spatial Transcriptomics Improves Cell Type Annotation and Reveals Critical Molecular Pathways"

### A Supplementary for robustness of Focus

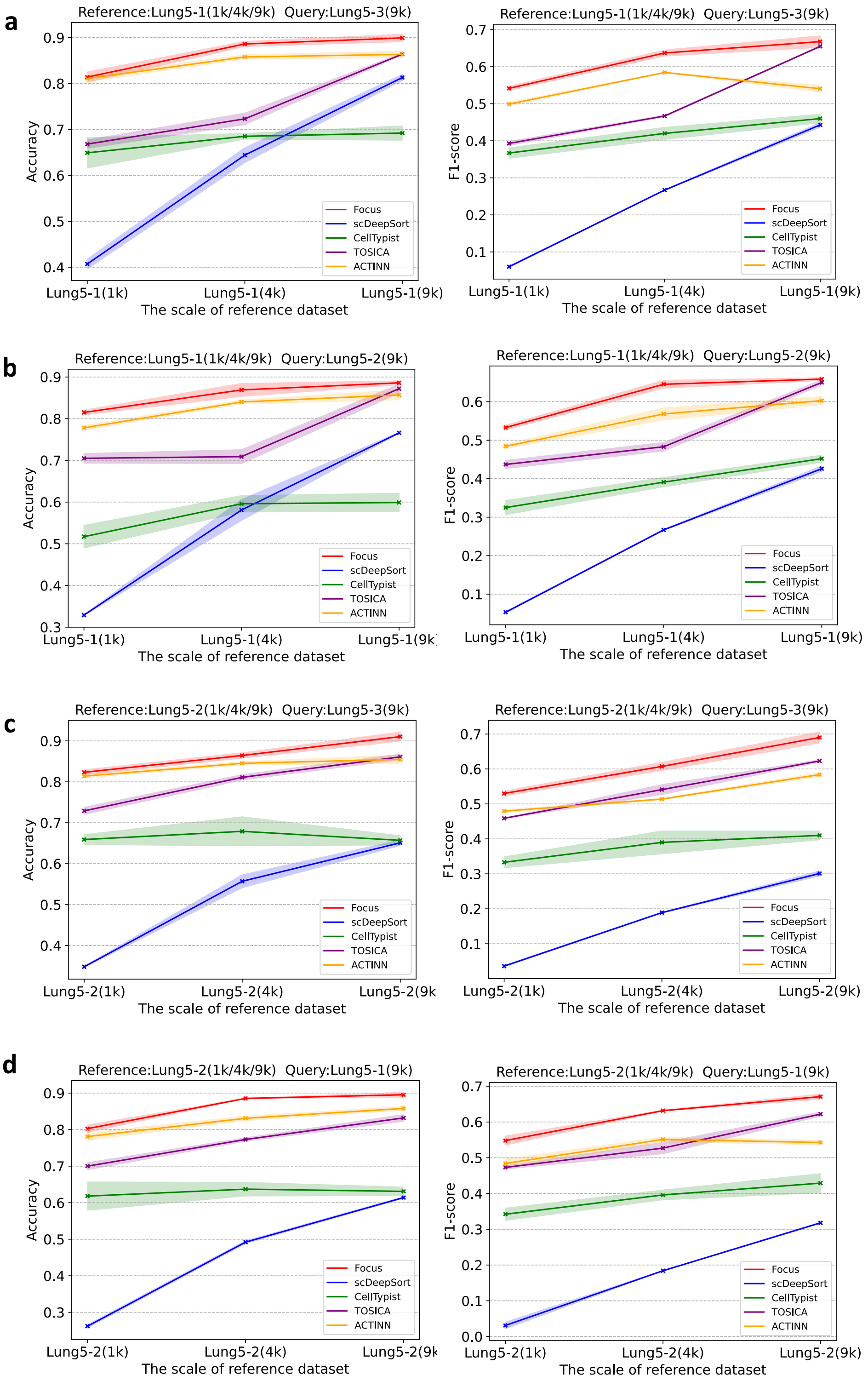


Figure 5: **Robustness of Focus and other methods to the reference data size.** **a**. Reference data: CosMx Lung5-1, Query data: CosMx Lung5-3. **b**. Reference data: CosMx Lung5-1, Query data: CosMx Lung5-2. **c**. Reference data: CosMx Lung5-2, Query data: CosMx Lung5-3. **d**. Reference data: CosMx Lung5-2, Query data: CosMx Lung5-1.

### B Supplementary of experimental setting

In our experiments, we selected 9,000 cells from various samples to evaluate the performance of our methods. When the reference and query datasets are from the same subject, we conducted experiments in the following settings: In the CosMx-Lung setting, we trained our models on CosMx Lung 5-2 and tested them on CosMx Lung 5-3. In the CosMx-Kidney setting, our training data came from CosMx Kidney 10838, and testing was performed on CosMx Kidney 1098. For the MERFISH-Mop setting, training was carried out using MERFISH mouse2_sample1, and testing occurred on MERFISH mouse2_sample2. In the Xenium-DCIS setting, we trained on Xenium replicate 1 and tested on replicate 2. However, when the reference and query datasets were obtained from different subjects, we conducted experiments in the following settings: In the CosMx-Lung setting, we trained on CosMx Lung 12 and tested on CosMx Lung 5-2. In the CosMx-Kidney setting, our training data came from CosMx Kidney 3323, and testing was performed on CosMx Kidney 10838. For the MERFISH-Mop setting, training was carried out using MERFISH mouse1_sample2, and testing occurred on MERFISH mouse2_sample2. These settings allowed us to assess the performance of our methods under various conditions and subject combinations.

### C Additional benchmarking results

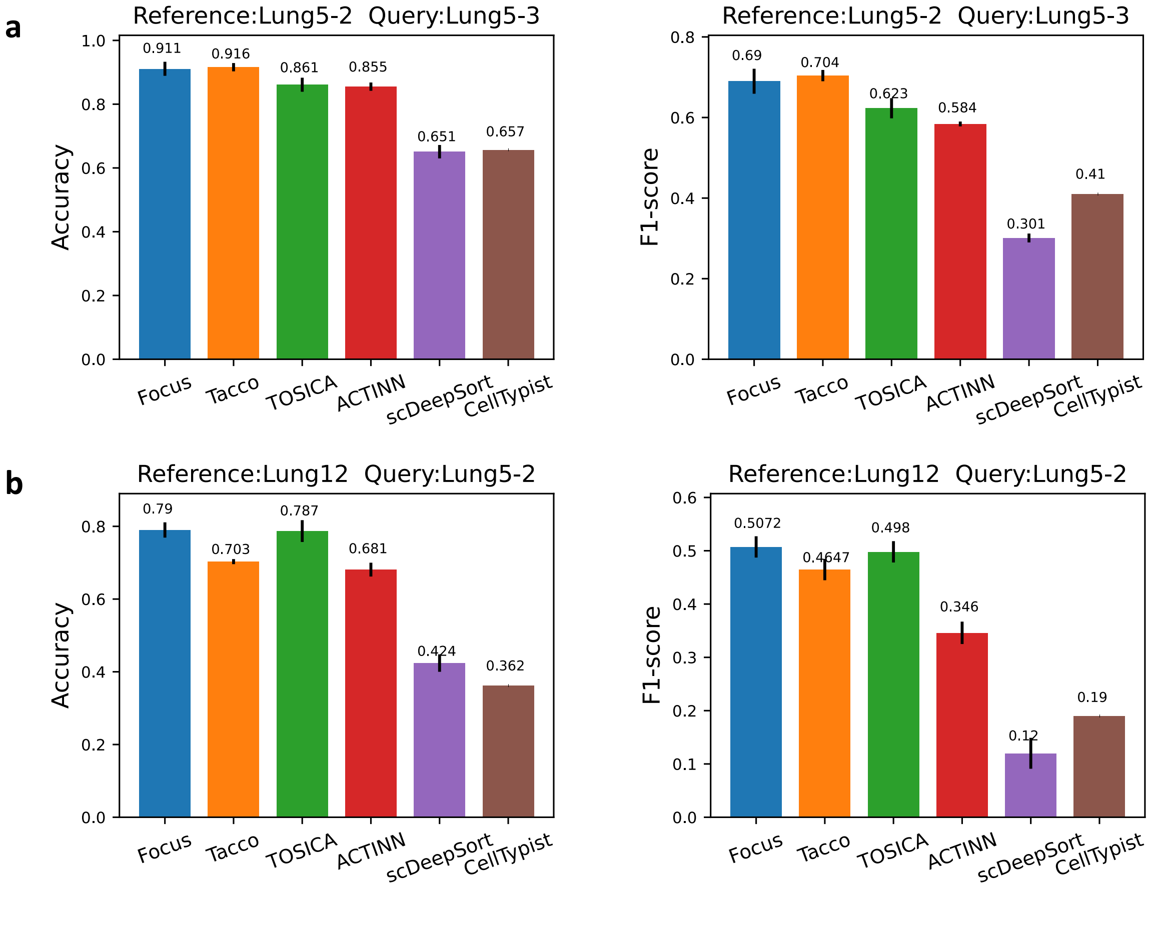


Figure 6: **Results of Focus and other methods on CosMx Lung dataset.** **a**. Reference data: CosMx Lung5-2, Query data: CosMx Lung5-3. **b**. Reference data: CosMx Lung12, Query data: CosMx Lung5-3.

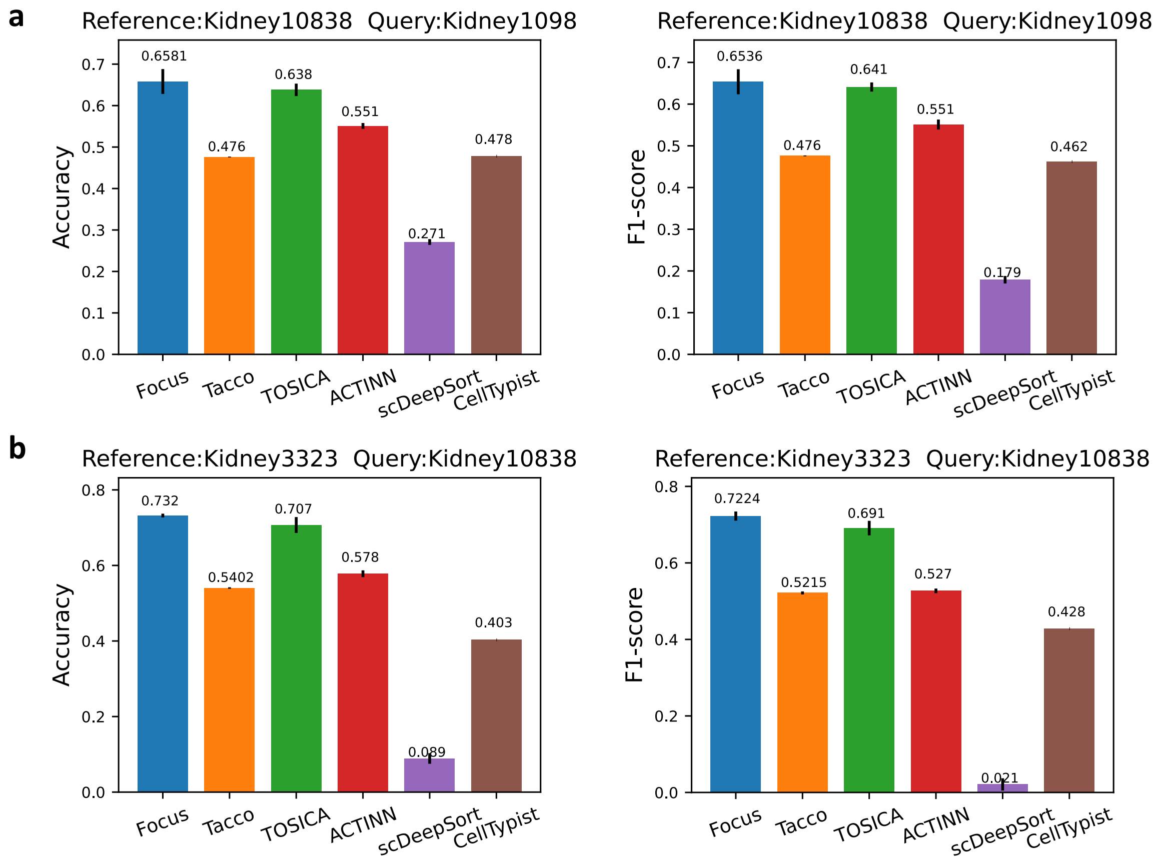


Figure 7: **Results of Focus and other methods on CosMx Kidney dataset. a**. Reference data: CosMx Kidney10838, Query data: CosMx Kidney1098. **b**. Reference data: CosMx Kidney3323, Query data: CosMx Kidney10838.

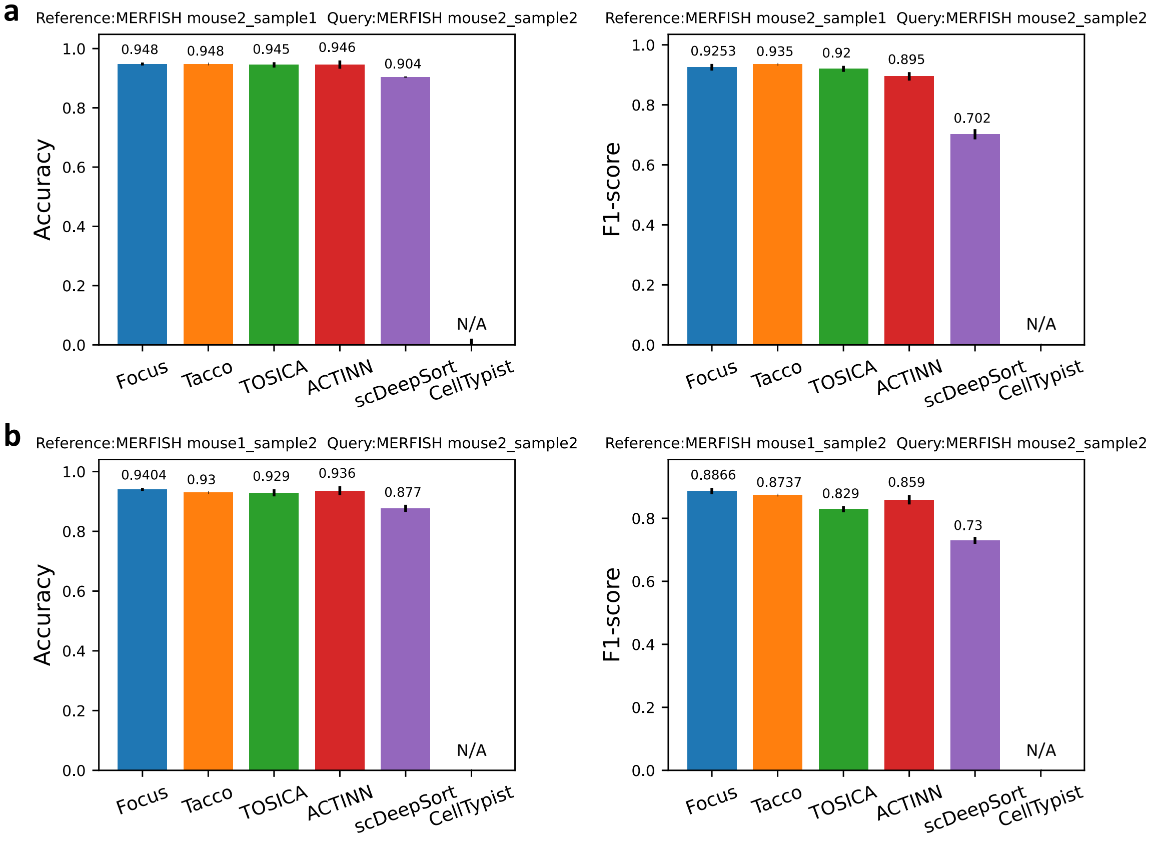


Figure 8: Results of Focus and other methods on MERFISH MOp dataset. a. Reference data: MERFISH mouse2_sample1, Query data: MERFISH mouse2_sample2. b. Reference data: MERFISH mouse1_sample2, Query data: MERFISH mouse2_sample2.

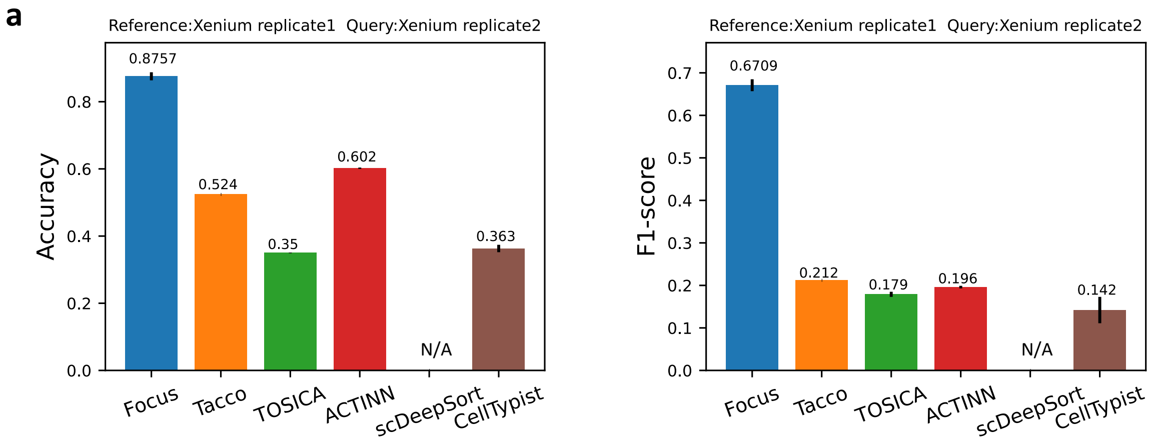


Figure 9: **Results of Focus and other methods on Xenium DCIS dataset. a**. Reference data: Xenium replicate1, Query data: Xenium replicate2.
